# The Leading-Edge Vortex of Swift Wings

**DOI:** 10.1101/099713

**Authors:** Rowan Eveline Muir, Ignazio Maria Viola

**Keywords:** bird wing aerodynamics, common swift *Apus apus*, dual leading-edge vortex, swept wing, delta wing, particle image velocimetry

## Abstract

Recent investigations on the aerodynamics of natural fliers have illuminated the significance of the Leading-Edge Vortex (LEV) for lift generation in a variety of flight conditions. A well documented example of an LEV is that generated by aircraft with highly swept, delta shaped wings. While the wing aerodynamics of a manoeuvring aircraft, a bird gliding and a bird in flapping flight vary significantly, it is believed that this existing knowledge will serve to add understanding to the complex aerodynamics of natural fliers. In this investigation, the wing of a common swift *Apus apus* is simplified to a model with swept wings and a sharp leading-edge, making it readily comparable to a model delta shaped wing of the same leading-edge geometry. Particle image velocimetry provides an understanding of the effect of the tapering swift wing on LEV development and stability, compared with the delta wing model. For the first time a dual LEV is recorded on a swift shaped wing, where it is found across all tested conditions. It is shown that the span-wise location of LEV breakdown is governed by the local chord rather than Reynolds number or angle of attack. These findings suggest that the common swift is able to generate a dual LEV while gliding, potentially delaying vortex breakdown by exploiting other features non explored here, such as wing twist and flexibility. It is further suggested that the vortex system could be used to damp loading fluctuations, reducing energy expenditure, rather than for lift augmentation.

---

> A dual leading-edge vortex is recorded on a swift shaped wing, suggesting that the common swift exploits this flow feature while gliding.

## 3 Introduction

The Leading-Edge Vortex (LEV) is a commonly found mechanism that under the correct conditions, can significantly augment the lift generation of both man-made and natural fliers (Ellington, 1999; Srygley and Thomas, 2002; Garmann *et al*., 2013; Jardin and David, 2014). The LEV is robust to kinematic change (Ellington, 1999) and has been identified across a wide range of Reynolds numbers (*Re*), from the laminar flow conditions (10 < *Re* < 10^4^) of autorotating seed pods (Lentink *et al*., 2009) and in insect (Ellington, 1999), bat (Muijres *et al*., 2008) and small bird (Lentink *et al*., 2007) flight, to the transitional and turbulent conditions of larger bird wings (Hubel and Tropea, 2010), fish fins (Borazjani and Daghooghi, 2013), delta wings (Gursul *et al*., 2005 and 2007), helicopter rotors (Corke and Thomas, 2015), sailing yachts (Viola *et al*., 2014) and wind turbines (Larsen *et al*., 2007). Increasingly driven by the potential exploitation of this effective lift mechanism for Micro Air Vehicles (MAV), and facilitated by improvements in flow visualisation and computational techniques, biologists and aerodynamicists now seek to further understanding of the flight of natural fliers at low to medium *Re* (10 < *Re* < 10^5^), as the LEV is increasingly understood to be a valuable flight mechanism to apply by design.

Across the broad range of flight conditions of natural fliers and aircraft, an LEV’s stability and ability to augment lift depend on numerous factors. These factors typically apply across the range, providing the potential for comparison, and include wing shape, sweep back angle (Λ), kinematics, and angle of attack (*α*). As an example, vortex lift is recorded at *α* = 30° on both the extensively studied slender delta wing (e.g. Visbal, 1996, where Λ = 75°) and the more complex, unsteady, flapping bird wing (Hubel and Tropea, 2010, where a blunt nosed model wing is tested at Λ = 10°). Both the wing kinematics, and the modelling or testing of natural flight, provide a significant challenge to researchers. Drawing analogy between the intricate LEV system of a natural flier and that of a more readily examined delta wing is a natural, though often difficult to undertake, starting point. This analogy has in fact been identified as critical in enhancing understanding of natural flight (e.g. Garmann and Visbal, 2014), and so may in turn be key in providing the necessary knowledge to allow this robust, high lift mechanism to be applied as required, by design.

Ellington *et al*., (1996) first noted the similarity in vortex structures, observing strong spiral vortices with notable axial velocity on the flapping wings of the hawkmoth *Manduca sexta* at 10^3^ < *Re* < 10^4^. They highlighted the resemblance to the spiral vortices generated over delta wings at significantly higher *Re*, while Usherwood and Ellington (2002) confirmed significant lift augmentation measuring a *C_l_* of 1.75 on a revolving (which replicates flapping without the supination and pronation movement) model hawkmoth wing. At insect scale the LEV is used at high angles of attack by butterflies, to generate ‘extreme’ lift (Srygley and Thomas, 2002), such as that needed during take off. The study supports the opinion of Ellington (2006) who notes that the high *α* of 30° – 40° required to generate an LEV naturally provides a highly unfavourable lift-to-drag ratio. He suggests that the rare gliding insects, damselfly *Calopteryx splendens* and dragonfly *Sympetrum sanguineum*, who have impressive lift-to-drag ratios of 6 to 21 at *Re* < 2 400, must seek to avoid leading-edge separation, and therefore the LEV, when gliding.

Conversely, Videler *et al*., (2004) show experimentally that the wing of a model common swift *Apus apus* is likely to generate an LEV while gliding; a result that may more conveniently be compared with the aerodynamics of a delta shaped wing. While similarly swept, the trailing edge shape and surface area of a swift wing differs from a delta shaped wing, tapering to a point from the body to the tip. The narrow, protruding, anterior vein of the primary feather however provides a sharp leading-edge that promotes separation, as is typical in delta wing design. With the model wings at a characteristic delta wing sweep Λ = 60°, flow separation is recorded as would be expected on a slender delta wing under the same conditions, and a single LEV is noted at low *α* (5° — 10°).

No force measurements are undertaken, but their assertion that the LEV is used by the swift for lift augmentation is challenged by Lentink *et al*. (2007) who test real, inherently flexible, wings with a maximum sweep of 50°; a more representative wing position during normal, non-diving gliding. They again find a single LEV but under different conditions; only at higher *α*, where *α* ≥ 11° and 14° at *Re* = 25 000 and 50 000, respectively. A maximum lift coefficient *C_l_* ≈ 0.9 at *α* = 30° is recorded from –6° ≤ *α* ≤ 30°. A fascinating study by Henningsson and Hedenström (2011) on live swifts, gliding in a wind tunnel, provides a calculated lift-to-drag ratio of between 9 and 12.5 across a *Re* range of 18 400 - 29 500, with a calculated maximum *C_l_* = 0.96. It is noted that this aligns well with the results of Lentink *et al*. (2007), although the maximum *α* attainable in the study is 6.3°. As Ellington (2006) commented regarding gliding insect flight, the impressive lift-to-drag ratio may suggest the lack of an LEV. The authors however note that while there is nothing to support its existence, there is nothing to discount it, either.

These discussions suggest both that swifts do not generate an LEV in gliding flight, where *α* is low, and that when they do, such as when manoeuvring and at high *α*, the LEV may not in fact be a high lift mechanism. This conclusion is interesting when compared with existing literature on the LEV, such as the lift coefficients presented on rigid and flexible delta wings of the same ‘low sweep’ Λ = 50° by Taylor *et al*. (2007). They show that at *α* = 30° the flexible wing benefits from *C_l_* ≤ 1.13, compared with a geometrically identical rigid wing generating *C_l_* ≤ 0.92 at the same *α*, though the tested *Re* is an order of magnitude higher than that of Lentink *et al*. (2007). Unfortunately a comparison of the flexibility of the real swift wing and the model delta wing cannot be made, and detail on leading-edge geometry is not reported by either authors.

A further comparison of interest is the investigation by Taylor *et al*. (2003), again of delta wing at Λ = 50° and low *Re*. They detail the development of not a single LEV, but a dual LEV, found at *Re* = 13 000, from 2.5° ≤*α* ≤ 15°. A dual LEV is a system commonly described on low sweep delta wings though to date, it is not known to have been identified on bird wings. It contains a larger, primary leading-edge vortex along with a second, minor, co-rotating vortex, separated by a region of counter rotating vorticity between the vortices and the wing suction surface. They further show that at *α* = 7.5°, the dual LEV is recorded across a range of *Re* overlapping the results of Lentink *et al*. (2007), from 8 700 to 34 700. Gordnier and Visbal (2005) similarly show the existence of a dual LEV computationally. At *Re* = 20 000 and 50 000, where *α* = 5° at the same Λ of 50°, both provide contrasting results to the ostensibly comparable experiments on a swift wing. It is not clear in either case what should cause this difference.

A low sweep of 35° ≤ Λ ≤ 55° is commonly seen in MAV wing design (Taylor *et al*., 2007), and understanding the particular geometry, wing construction, and set of conditions required to optimise lift and manoeuvrability at lower *Re* is of particular interest to the research community. Many seek to learn from the skilled and precise aerial acrobatics of birds in flight, however is would appear that significant aerodynamic differences exist, even under seemingly comparable test conditions. Neither an LEV at low *α* (≤ 11°), nor a dual LEV, has been identified on the swift wings discussed, in comparison with delta wing research which notes the existence of a dual LEV even at low *α* (2.5°).

To better compare the aerodynamics of the two discussed, we perform flow measurements at *Re* typical of the gliding swift, between 12 000 and 67 000 (Lentink and de Kat, 2014), over a model swift wing planform and a model delta shaped wing of readily comparable geometry. This allows the isolation of the effect of the swift wing trailing edge shape, and the form and evolution of the LEV across a range of *α* and *Re* is reviewed for both planforms. The aim of this investigation is to identify whether a swift shaped wing can generate a comparable LEV system to that of a delta wing, to explore the effect of the trailing edge shape of the swift wing on LEV generation and stability, and to discuss the results in relation to known swift flight aerodynamics.

## 4 Materials and Methods

### 4.1 Model specification and facility

The tested non-slender delta wing (Figure 1) has Λ = 50°, root chord *c_r_* = 0.15 m and wing span *b* = 0.25 m, which with area *S* gives the wing an aspect ratio (*AR* = *b*^2^/*S*) of 3.36. This design was then replicated and modified to produce a simple, rigid, swift shaped wing planform based on the geometry provided in Lentink *et al*. (2007); the wing shape only is constructed, without the body, with Λ = 50°, b = 6*c_r_* and *AR* = 6. While that paper and others discuss the importance of wing flexibility (e.g. Shyy *et al*., 2010; Cleaver *et al*., 2014) and the spanwise variation in angle of attack common in insect and bird wings due to twist (e.g. Hubel and Tropea, 2010), no research could be found on the effect of wing geometry alone in the two readily comparable cases of a swift shaped wing and a low sweep delta wing. The rigid design will therefore allow new insights on the fundamental differences (and similarities) between the two geometries.

**Figure 1:**
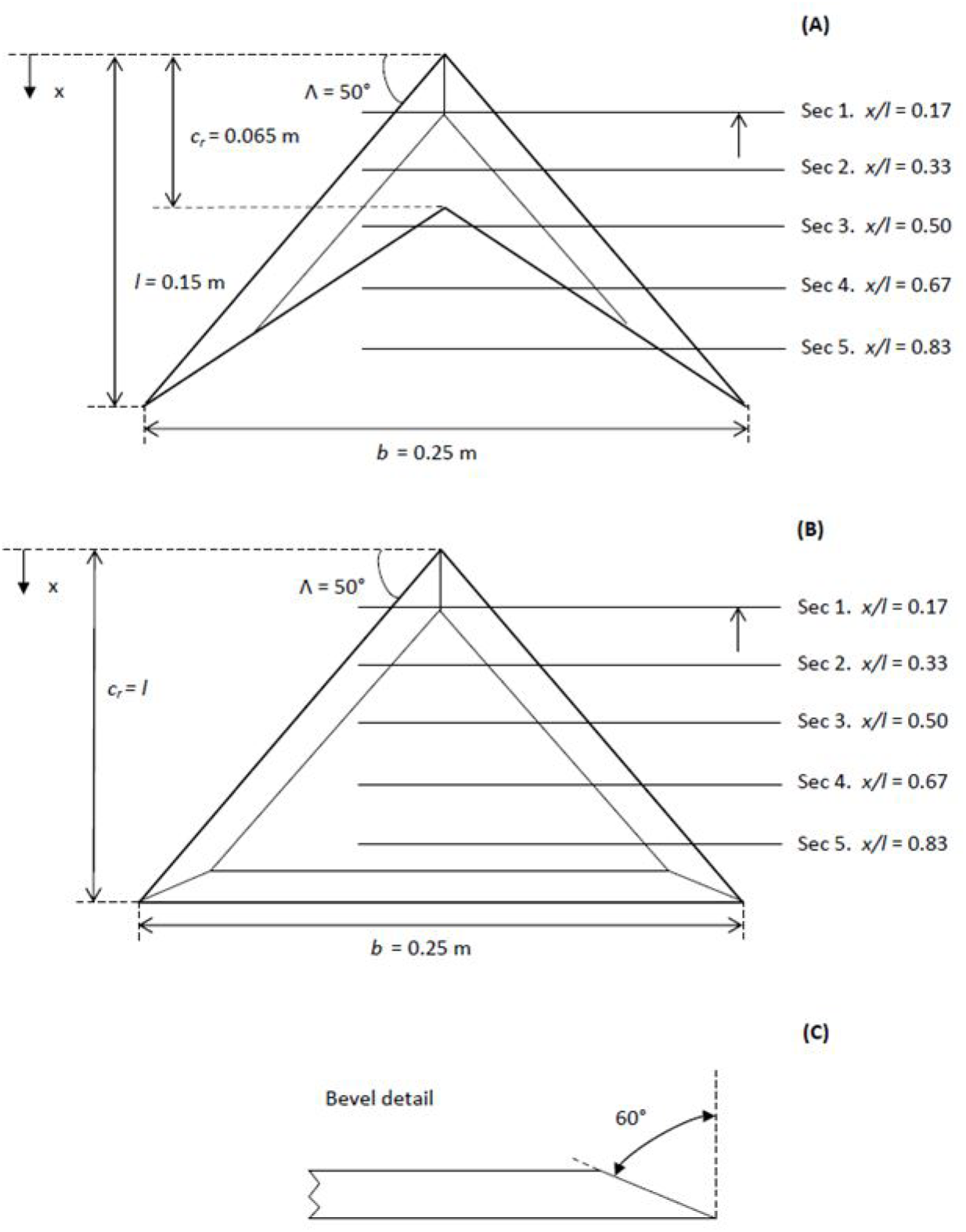
Schematic diagram of the tested models. (A) swift shaped wing, (B) delta shaped wing, and (C) bevel detail.

The wing sections were laser cut from 10 mm thick acrylic. A sharp leading-edge is provided on both wings by applying a 60° bevel on the windward side, as is common practice in low sweep delta wing investigations (e.g. Ol and Gharib, 2003; Wang *et al*., 2007). The leading-edge bevel also results in the wing gaining an effective camber (Gursul *et al*., 2005; Verhaagen, 2012). Camber can however be seen on the cross section of bird wings (eg Videler *et al*., 2004), and as aerodynamic forces are not explored in this instance this was not thought to be unduly detrimental to the study. The effect of the edge bevel on the flow field is discussed in the results.

The wing was rigidly located in a low speed, free surface, water flume at the University of Edinburgh (Figure 2). The glass walled flume is 8 m long, 0.4 m wide and the water depth was set to 0.5 m; it is of closed circuit recirculating current design, and includes a series of meshes around 4 m upstream of the model to reduce turbulence. The water speed is controlled by an electric motor driving a propeller providing a flow speed (U_0_) at the model location of up to 1 ms^−1^. In the present investigation 0.10 < *U*_0_ < 0.44 ms^−1^ resulting in a chord based *Re* range of around 15 800 to 65 500, with turbulence ranging from 6.8% to 3.1% respectively.

**Figure 2:**
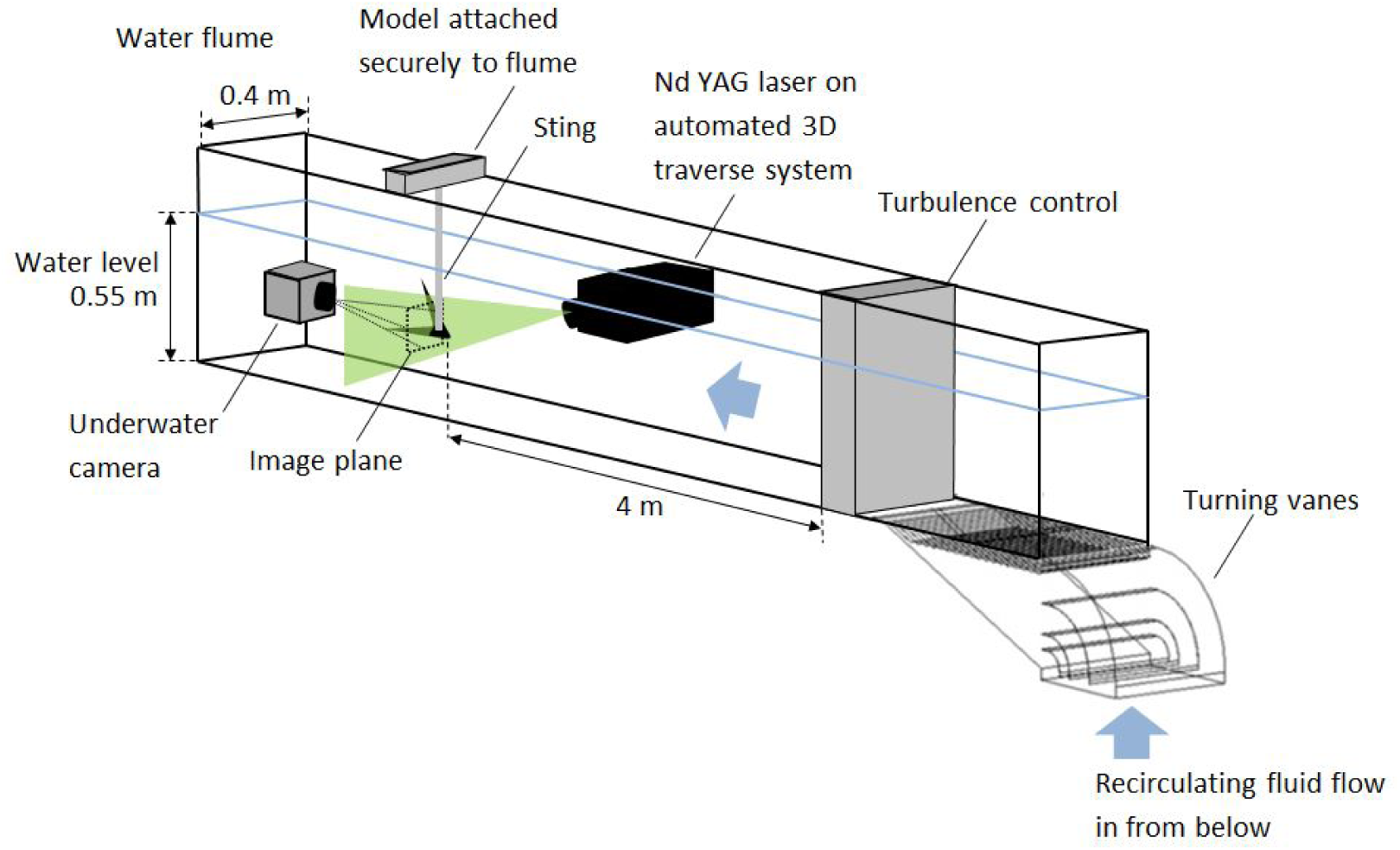
Schematic diagram of the experimental rig. The figure shows the general position of the laser, the underwater camera and the model in the flume.

**Figure 3:**
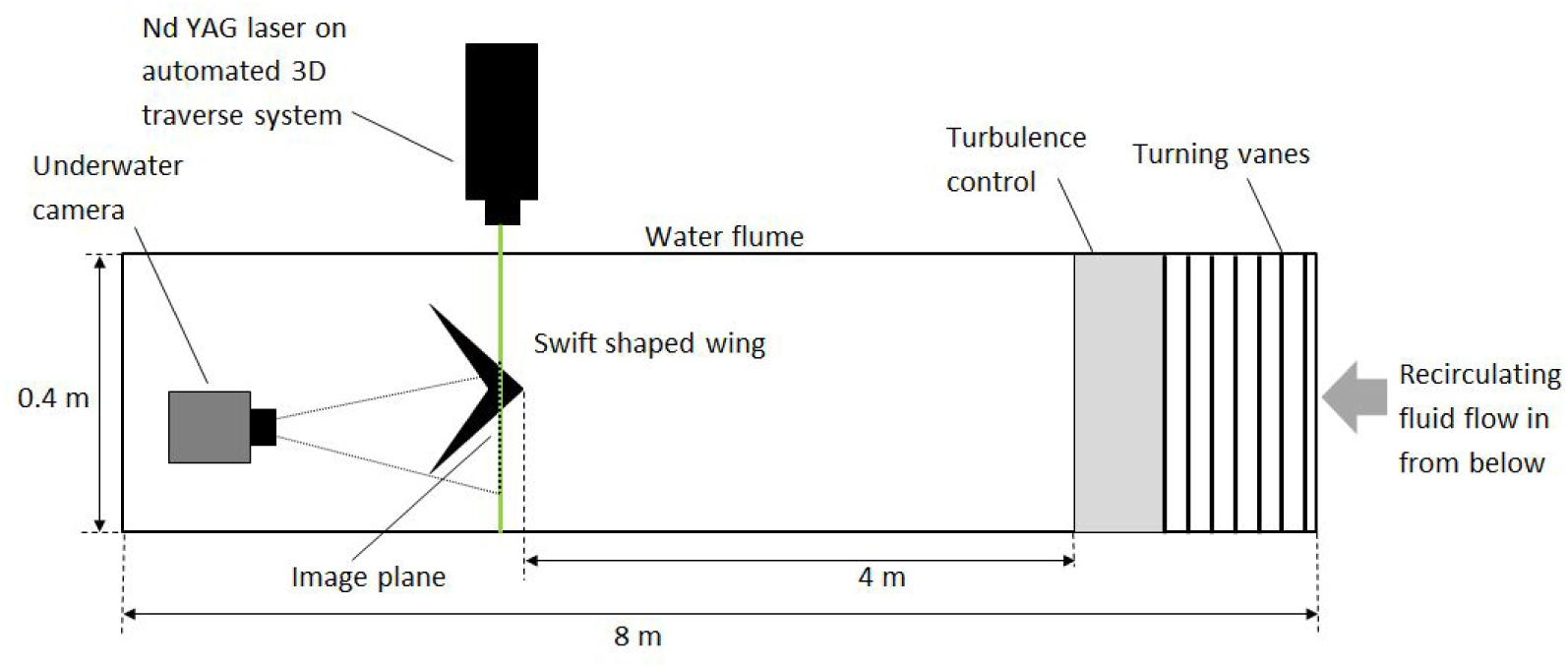
Plan view schematic of the experimental rig.

The models were supported by a frame securely mounted on the top of the flume, and via a 12 mm diameter cylindrical sting attached to the upper, bevelled surface of the wing (Figure 2). The wing was attached with a screw countersunk into, and flush with, the suction surface; the suction surface was therefore the lower surface of the wing. The sting was attached to the frame via an adjustable bracket that allowed the angle of attack of the wing to be modified. The angle was measured and set by a digital level with accuracy of ±0.01°.

A set of images were taken at *x*/*l* = 0.33, where x is a chordwise coordinate from the apex of the wing and *l* is the chordwise length of the wing. This location was selected to be sufficiently far from the apex to allow development of the flow feature, however upstream of vortex breakdown in most cases. The flow field was recorded at the angle at which a vortex was identified by Lentink *et al*. (2007), *α* = 15°, then again at steps reducing the angle of attack to *α* = 0°. While more *α* values were tested, only *α* = 0°, 5°, 10° and 15° are presented. A set of images were also taken at *α* = 7.5°, a previously documented flow regime on non-slender delta wings to allow an element of validation (e.g. Taylor *et al*., 2003), at *x*/*l* = 0.5 on both the swift and delta shaped wings. The first and second set of images were taken across the range of *Re* however only the lowest (*Re* = 15 800) and highest (*Re* = 65 500) cases are presented. A third set of images were again taken at *α* = 7.5°, but at all 5 locations along the length of each wing at *Re* = 17 500 to elucidate vortex development along the leading-edge of each planform more fully.

### 4.2 Particle Imaging Velocimetry

A dual-pulse Nd:YAG laser (15 - 200 mJ at 532 nm, 200 Hz) was used to generate a 3 mm thick light sheet illuminating seeding particles in selected planes at each *α*. The seeding particles used are Conduct-O-Fil silvered spheres, with an average diameter of 14 *μ*m and average density of 1.7 g/cc. A LaVision Imager pro SX 5M camera fitted in a waterproof housing was secured in the flume downstream of the wing support rig. In order to improve resolution, only one half of the swift or delta wing was imaged. All images are available on the Edinburgh DataShare repository (http://datashare.is.ed.ac.uk) and on Dryad (http://datadryad.org).

Analysis was undertaken using the LaVision Particle Image Velocimetry (PIV) software, DaVis 8.3.0. The field of view of around 150 × 125 mm was broken into interrogation windows and analysed using a multi pass cross correlation algorithm. A primary pass was carried out with a window size of 64 × 64 pix and a 50% overlap, which was then refined with 3 passes at 32 × 32 pix window size and a 75% overlap. Outliers were removed in post-processing with a median filter, and a 3 × 3 Gaussian smoothing was applied. Flow features were found to be steady; unless otherwise noted, results presented are averages of 100 image pairs taken at a frequency of 7.5 Hz with a resolution of 2448 × 2050 pixels.

## 5 Results

### 5.1 Primary and secondary LEV

Both a primary and second minor LEV was found at all *α* and *Re* tested on both wings, showing for the first time that a dual LEV can exist on a swift shaped wing. Figure 4 shows the in-plane vector velocity field and the vorticity contours (non-dimensionalised by *U*_0_ and *c_r_* = 0.15 m) of the flow over the swift shaped wing at a position of *x*/*l* = 0.33. The wing appears to generate a coherent, stable dual LEV system in all cases examined; from *α* = 0° to 15° and *Re* = 15 800 to 65 500. Vortex breakdown is known to move upstream with increasing *α* (e.g. Ol and Gharib, 2003); in the present investigation breakdown dominates the flow by *α* = 15° and is imaged. As seen in similar investigations on delta wings (e.g. Taylor *et al*., 2003) the broken down vortex remains located over the wing, with the vertical position of the vortex axis largely unaffected.

**Figure 4:**
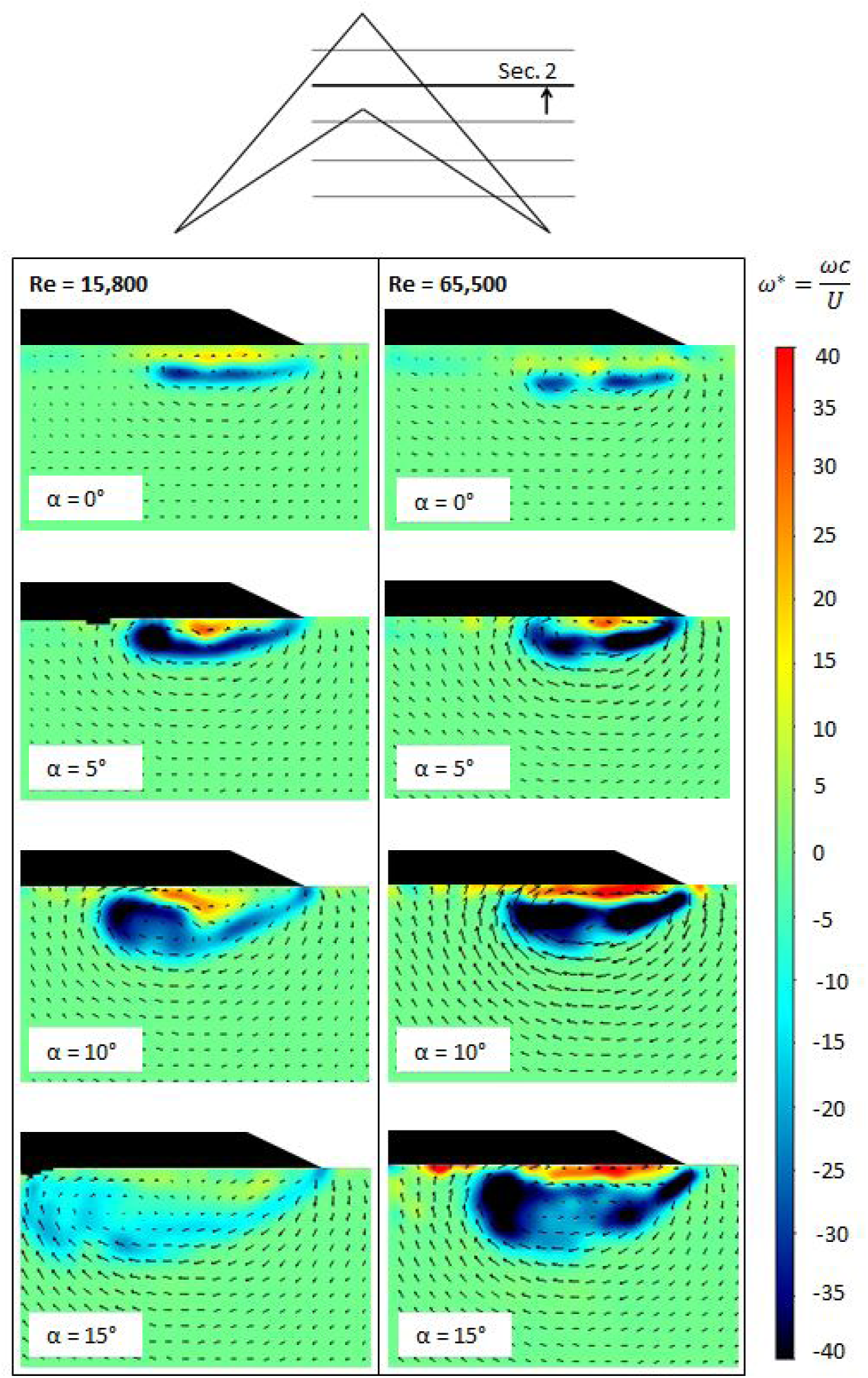
In-plane velocity vectors and vorticity contours around the swift shaped wing. Tests performed at *Re* = 15 800 and 65 500 for a range of *α*.

The identification of a dual LEV in all cases tested is contrary to the previous PIV results of Videler *et al*. (2004) and the tuft flow visualisation of Lentink *et al*. (2007). In the present study, flow separates from the leading-edge forming a strong shear layer, which then rolls up into a coherent LEV. In line with previously reported low sweep delta wing aerodynamics (e.g. Ol and Gharib, 2003; Taylor *et al*., 2003; Gursul *et al*., 2005), secondary separation of opposite sign can be seen between the LEV and the wing surface, formed by the interaction between the LEV and the boundary layer. Dual vortices more readily develop on low sweep delta wings due to the increasing proximity of the LEV to the wing suction surface with reducing sweep (Gursul *et al*., 2005). The two distinct co-rotating vortices, one larger primary vortex and one minor vortex, inboard and outboard of the secondary separation, respectively, are clearly seen in Figure 4, most distinctly at *α* = 10° and *Re* = 15 800.

The existence of an LEV at zero angle of attack results from an effective camber introduced by the leading-edge bevel; the zero lift angle was found to be around *α* = –4°, resulting in a weak dual LEV structure being visible at *α* = 0°. As reported in previous investigations on both delta and flapping/rotating wings, (e.g. Ozen and Rockwell, 2011; Wojcik and Buchholz, 2014; Gursul *et al*., 2005), the LEV system increases in size and strength with *α* resulting in a reduced proximity to the wing surface and to the leading-edge. With increasing *Re* the vortex again responds as expected, becoming stronger and more concentrated, and moving outboard with the reduction in viscous effects (Taylor *et al*., 2003). Most notably this is demonstrated at the highest *α* where the lower *Re* case displays vortex breakdown.

### 5.2 Comparison between swift shaped wing and delta wing vortex structures

Figure 5 shows that under the same hydrodynamic conditions, the flow around the delta and swift shaped wings is highly comparable, both developing a coherent dual vortex flow structure located around the same horizontal and vertical position over the suction surface of the wing. The primary difference appears to be the increased size, but reduced coherence and strength, of the swift vortex structure at the lower *Re*, compared with that of the delta wing. The strength and magnitude of the secondary vortex in the swift wing case is also reduced. At higher Re the difference between the two systems is reduced but still may be perceived. As previously described for Figure 4, the delta wing vortex structure shifts position slightly as expected (Taylor *et al*., 2003) with increasing *Re*.

**Figure 5:**
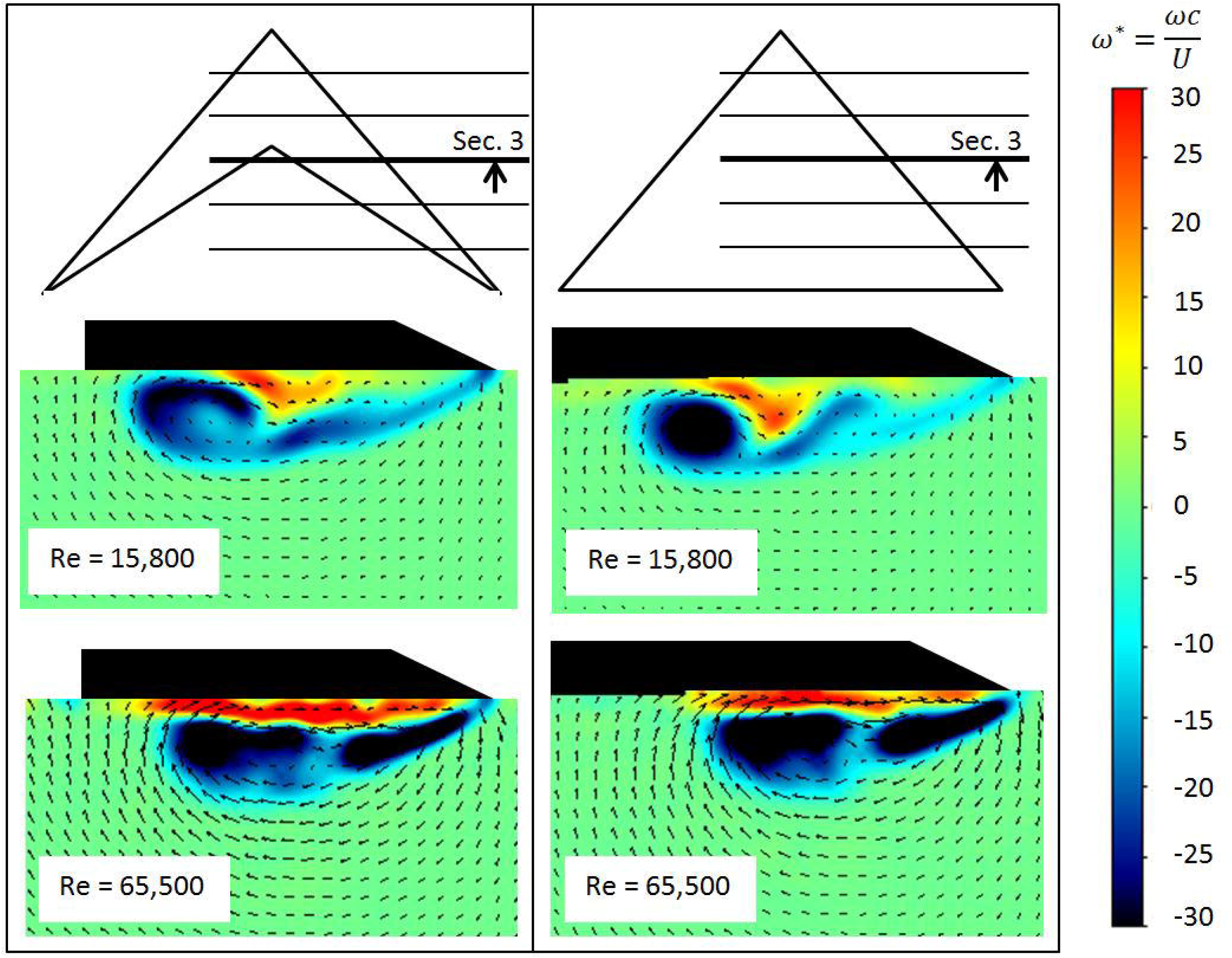
In-plane velocity vectors and vorticity contours around the swift and delta shaped wings. Tests performed at *α* = 7.5° and *Re* = 15 800 and 65 500.

### 5.3 Vortex breakdown

Figure 6 shows the vortex development along the span of the swift and delta wings at *α* = 7.5° and *Re* = 17 500. While the flow towards the root of the wing is highly comparable in vorticity magnitude, location and coherence, differences are seen from around *x*/*l* = 0.5 (Sec. 3). In the case of the swift shaped wing, the vortex system suffers breakdown as its size exceeds that of the local wing chord beneath it and flow re-attachment is no longer possible. Instead, an area of separated flow is seen beneath the still obvious shear layer. Despite breakdown, the shear layer and primary LEV still display two distinct areas of vorticity. While the effect of disorder on the averaged image of the flow is to give the appearance of largely reduced vorticity, instantaneous images show that high vorticity in the region remains, supporting existing research (e.g. Gursul *et al*., 2005) showing that a broken down vortex continues to provide lift benefits. The dual vortex system on the delta shaped wing begins to show signs of breakdown only in the last image, where the second minor vortex displays the greatest effect. This is in line with the experimental results of, for example, Taylor *et al*. (2003) and Ol and Gharib (2003). It is clear that the vortex system of the delta shaped wing has improved stability and coherence towards the wing tip as it continues to be supported by the suction surface of the wing, and the flow is able to reattach inboard of the vortex system.

**Figure 6:**
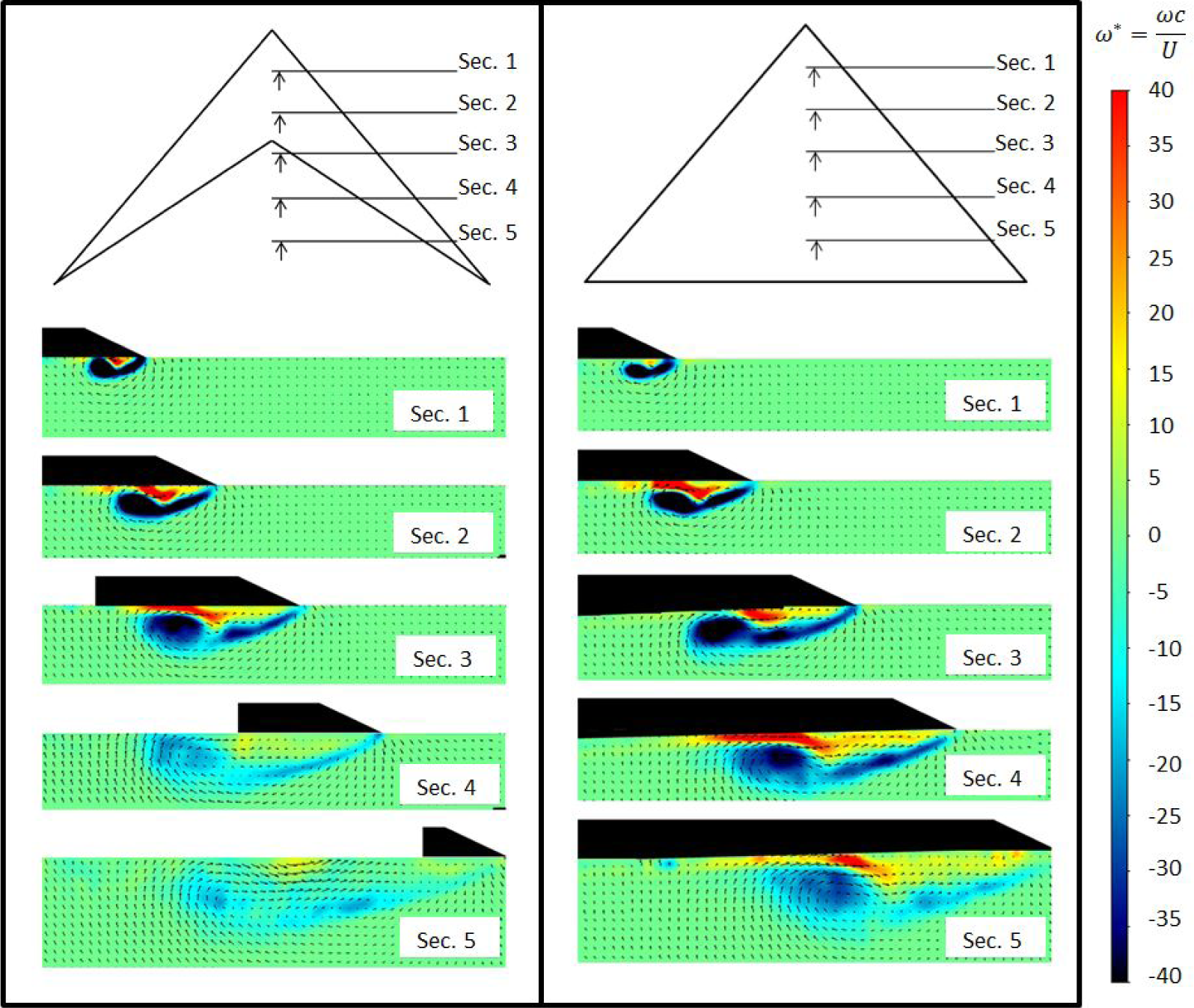
In-plane velocity vectors and vorticity contours around the swift and delta shaped wings. Tests performed at *Re* = 17,500 and *α* = 7.5°.

## 6 Discussion

The images presented show for the first time that a swift shaped wing can generate a stable and coherent dual LEV system, in line with that found on a delta wing of comparable geometry, but under typical gliding conditions of a swift; at low *α* and *Re*.

While the PIV measurements of Videler *et al*. (2007) found a prominent but single LEV above the hand wing section on a swift wing where *α* > 5°, their model had a slender sweep of 60°, bringing improved stability and coherence of the LEV. Reducing Λ to 50° as in the present work would increase the minimum *α* required for LEV stability from the 5° found, providing increased contrast to those results presented now. Similarly, Lentink *et al*. (2007) tested real swift wings at Λ = 50° and also did not record an LEV at low angles of attack, finding them only at *α* ≤11° and *α* ≤ 14°. No dual vortex was recorded in either case.

A comparison of these results appears to suggest that the increased *α* required to generate an LEV on the swift wings tested previously is not due to the wing trailing edge geometry, but to another flow condition or design parameter. It is easily conceivable that the differences in results noted here are due to varying sharpness of the leading-edge or wing thickness across the respective models tested. In the case of the dual LEV, the surface texture or structure of the bird wing could affect the nature of the secondary separation and its influence on the vortex structure.

It may also be that the flow visualisation method used is key in identifying the smaller, weaker vortex structure at low *α* and Λ. Lentink *et al*. (2007) use a tuft grid survey, originally developed for testing flow over slow flying delta wings (Bird, 1969). The tuft grid method places a tuft (or single hair) at regular intervals across the wing which can then be imaged as it moves according to the local flow regime. Where the size, strength, position *e.t.c* of the flow feature is not known, it can be difficult to suitably define the length, stiffness and position of the tuft to highlight the sought feature. It may be possible that a weak LEV did exist in the cases presented, but that the flow visualisation technique used was not sensitive enough to capture it. Similarly, a dual vortex may have existed at higher angles of attack with the weaker minor vortex not identified for the same reason.

Figure 5 allows a comparison between the vortex system on the swift shaped wing and the delta wing. The swift shaped wing appears to generate slightly larger vortices than the delta wing, however they are less coherent, and contain a reduced area of concentrated vorticity. This reduction in coherence is associated with reduced lift, while the increase in size also increases drag, further reducing the lift-to-drag ratio and the potential benefit, to the bird in gliding flight, of the lift generated.

The size of the LEV provides an indication of the balance of vorticity being introduced to the vortex or vortex system from the separated shear layer, and the vorticity that is extracted from the same and passed down-stream. When insufficient vorticity is extracted, it is instead accumulated within the LEV, which grows in size until it is convected downstream and a new LEV is formed. This results in periodic shedding of vorticity. In the case of swept wings, vorticity is extracted primarily through two means. The greater the sweep angle of the wing, the larger the component of the velocity being affected in the spanwise direction, and thus the component of flow pushed towards the tip. The increasing vorticity toward the wing tip due to increased feeding from the separated shear layer results in a conical vortex such as that seen here, and an associated spanwise pressure gradient. The stability of the vortex and its generated lift thus largely depends on the wing geometry and flow conditions; where too low to balance the vorticity production periodic shedding occurs, and when vorticity is extracted at a higher rate, a small, highly coherent vortex can be generated. The observed increase in size of the swift wing vortex system, therefore, suggests reduced extraction when compared with the delta wing.

Figure 6 shows the swift wing vortex before and after the point of breakdown. Interestingly, the spanwise location of the breakdown of the vortex in the swift shaped wing appears to be driven by the size of the vortex relative to the local chord of the wing; where the fluid flowing over the LEV is no longer able to reattach inboard of the vortex due to tapering of the wing, the vortex becomes unsteady and breaks down. In the case presented, the breakdown of the swift wing vortex reduces the pressure gradient along the length of the remaining vortex upstream of the point of breakdown; the rate of vorticity extraction would therefore be lowered, increasing the size and reducing the coherence of the vortex system. While the swift wing shape does not prevent the generation of a vortex system at low *α*, it does provide a physical limit to any development of the LEV that may be caused by increasing *α*, the advance ratio or Re. Breakdown instead results, increasing drag. This is not a limit experienced by the delta shaped wing, and it is worthy of note, as it affects both the ability of the swift wing to generate lift, and its lift-to-drag ratio.

The effect of the swift wing geometry is exaggerated in the test case presented here by the more pronounced reduction in chord length of the model, compared with that of a typical swift wing planform (Henningsson and Hedenström, 2011). A twisted leading-edge and wing flexibility, as previously discussed, would also affect the breakdown location; a spanwise reduction in the effective angle of attack via twist or flexibility would attenuate vortex development towards the wing tip, delaying the onset of vortex breakdown.

The many differences between the wing of a real swift, such as that tested by Lentink *et al*. (2007), and the model presented here have arguably been ‘selected’ in natural fliers to improve their aerodynamic and flight ability. The leading-edge detail, in particular the sharpness, is known not only to promote flow separation, but also to define the relationship between *α* and the force coefficients generated (Usherwood and Ellington, 2002), while the behaviour of the LEV is increasingly dependant on leading-edge shape at low sweep (Miau *et al*., 1995). If a vortex can be generated at low *α*, promoted by the sharp leading edge of the swift wing and stabilised in part by leading edge twist and wing flexibility, it is interesting to consider what function it may serve if not one of lift augmentation, or lift-to-drag ratio improvement. While Henningsson *et al*. (2014) conclude that the swift wing is best adapted for flapping rather than gliding flight, the ability to generate and maintain a stable leading-edge vortex across a wide range of *α* may be a useful addition to the suite of flight optimisation tools deftly deployed by the swift. As their distinction *apodiformes* (the Greek for ‘footless’) eludes to, swifts spend the majority of their lives in the air; hunting, eating, collecting water, mating and even roosting (Lentink *et al*., 2007) and they have necessarily evolved to be one of the most aerodynamically refined bird species (Lentink and Kat, 2014). The optimisation of energy required to undertake these necessary activities is clearly essential, and with the swift spending a significant portion of its flight time gliding, often in complex air flows, any reduction in gliding energy requirement would be of significant benefit.

While the LEV is typically referred to as a high lift mechanism (Ellington, 1999; Srygley and Thomas, 2002; Lentink *et al*., 2009; Garmann *et al*., 2013; Jardin and David, 2014), it seems that this is the case only within a specific combination of conditions. It may be that the ability to generate an LEV and/or LEV system on a swift wing can provide another function. Along with potentially enhancing lift and manoeuvrability at high *α* and during flapping flight, the LEV system may help to manage wing loading and aerodynamic force fluctuations at lower α, by acting as a dampening mechanism. The ‘robustness to kinematic change’ noted by Ellington (2009) may allow the LEV to react more slowly to sudden changes in angle of attack that might otherwise have resulted in a sharp increase in lift, or in attached flow separating and the wing stalling. Rather, the LEV may grow or reduce in size at a rate mediated by inertia of the vortex system, altering the lift-to-drag ratio, but preventing against the severe loading fluctuations associated with stall. The ability to exploit the LEV in this way could save the bird significant expenditure of energy, and may be a useful addition to a suite of flow control techniques utilised by MAVs seeking to fly in complex or turbulent weather conditions.

## 7 Authors’ contributions

REM and IMV jointly conceived and designed the study. REM was responsible for the construction of the model, the execution of the experiments and the data analysis; she prepared the original draft of the manuscript; IMV was responsible for supervision and administration of the project, and funding acquisition; he contributed to the critical analysis and interpretation of the results; and reviewed and edited the manuscript. Both authors gave final approval for publication.

## 8 Acknowledgements

The authors would like to thank Susan Tully and Jean-Baptiste Richon for their experimental insight and guidance throughout.

## 2 List of Symbols and Abbreviations

*AR*: Aspect ratio (*b*^2^/*S*)
*b*: Span (m)
*c*: Local spanwise chord (m)
*c_r_*: Root chord (m)
*C_l_*: Lift coefficient
*l*: Chordwise length of the wing (m)
*R_e_*: Reynolds number
*S*: Area (m^2^)
*U*_0_: Free stream velocity (ms^−1^)
*x*: Chordwise coordinate (m)
*α*: Angle of attack (deg.)
Λ: Sweep back angle (deg.)
*ω*: Chordwise vorticity (s^−1^)
*ω*\*: Non-dimensional chordwise vorticity (*ωc*/*U*)
LEV: Leading-Edge Vortex
MAV: Micro air vehicle
PIV: Particle Image Velocimetry

## References

Bird, J.D. Tuft-grid surveys at low speeds for delta wings. NASA Technical Note D-5045 (1969).

Borazjani, I., and Daghooghi, M. (2013). The fish tail motion forms an attached leading edge vortex. Proceedings. Biological Sciences / The Royal Society, 280, 20122071. http://doi.org/10.1098/rspb.2012.2071

Cleaver, D. J., Gursul, I., Calderon, D. E., and Wang, Z. (2014). Thrust enhancement due to flexible trailing-edge of plunging foils. Journal of Fluids and Structures, 51, 401–412. doi:10.1016/j.jfluidstructs.2014.09.006

Corke, T. C., and Thomas, F. O. (2015). Dynamic Stall in Pitching Airfoils: Aerodynamic Damping and Compressibility Effects. Annual Review of Fluid Mechanics, 47, 479–505. doi: 10.1146/annurev-fluid-010814-013632

Dudley, R. (1991). Biomechanics of flight in neotropical butterflies: Aerodynamics and mechanical power requirements. J. Exp. Biol. 159, 335–357.

Ellington, C. P. (1984). The aerodynamics of flapping insect flight. Philosophical Transactions of the Royal Society of London [Biology], 24, 95–105.

Ellington, C. P. (1999). The novel aerodynamics of insect flight: applications to micro-air vehicles. The Journal of Experimental Biology, 202, 3439–3448.

Ellington, C. P. (2006). Insects versus birds: the great divide. In 35th AIAA aerospace science meeting and exhibit (pp. 450–455).

Ellington, C. P., Van Den Berg, C., Willmott, A. P., and Thomas, A. L. (1996). Leading-edge Vortices in Insect Flight. Nature, 384(19/26), 626–630.

Garmann, D. J., and Visbal, M. R. (2014). Dynamics of revolving wings for various aspect ratios. Journal of Fluid Mechanics, 748, 932–956. doi:10.1017/jfm.2014.212

Garmann, D. J., Visbal, M. R., and Orkwis, P. D. (2013). Threedimensional flow structure and aerodynamic loading on a revolving wing. Physics of Fluids, 25(3), 034101. doi:10.1063/1.4794753

Gordnier, R. E., and Visbal, M. R. (2005). Compact Difference Scheme Applied to Simulation of Low-Sweep Delta Wing Flow. AIAA Journal. doi:10.2514/1.5403

Gursul, I., Gordnier, R., and Visbal, M. (2005). Unsteady aerodynamics of nonslender delta wings. Progress in Aerospace Sciences, 41(7), 515–557. doi:10.1016/j.paerosci.2005.09.002

Gursul, I., Wang, Z., and Vardaki, E. (2007). Review of flow control mechanisms of leading-edge vortices. Progress in Aerospace Sciences, 43(7-8), 246–270. doi:10.1016/j.paerosci.2007.08.001

Henningsson, P., and Hedenström, A. (2011). Aerodynamics of gliding flight in common swifts. The Journal of Experimental Biology, 214(Pt 3), 382–393. doi:10.1242/jeb.050609

Henningsson, P., Hedenstrm, A., and Bomphrey, R. J. (2014). Efficiency of lift production in flapping and gliding flight of swifts. PLoS ONE, 9(2). http://doi.org/10.1371/journal.pone.0090170

Hubel, T. Y., and Tropea, C. (2010). The importance of leading edge vortices under simplified flapping flight conditions at the size scale of birds. The Journal of Experimental Biology, 213(11), 1930–9. doi:10.1242/jeb.040857

Jardin, T., and David, L. (2014). Spanwise gradients in flow speed help stabilize leading-edge vortices on revolving wings. Physical Review E, 90(1), 013011. doi:10.1103/PhysRevE.90.013011

Jin-Jun, W., and Wang, Z. (2008). Experimental Investigations on Leading-Edge Vortex Structures for Flow over Non-Slender Delta Wings. Chinese Physics Letters, 25(7), 2550–2553. doi:10.1088/0256-307X/25/7/060

Johansson, L. C., Engel, S., Kelber, A., Heerenbrink, M. K., and Hedenström, A. (2013). Multiple leading edge vortices of unexpected strength in freely flying hawkmoth. Scientific Reports, 3, 3264. doi:10.1038/srep03264

Larsen JW, Nielsen SRK, Krenk S. (2007). Dynamic stall model for wind tubine airfoils. Journal of Fluids and Structures, 23, 959–82. doi:10.1016/j.jfluidstructs.2007.

Lentink, D., and de Kat, R. (2014). Gliding swifts attain laminar flow over rough wings. PloS One, 9(6), e99901. doi:10.1371/journal.pone.0099901

Lentink, D., and Dickinson, M. H. (2009). Rotational accelerations stabilize leading edge vortices on revolving fly wings. The Journal of Experimental Biology, 212(Pt 16), 2705–19. doi:10.1242/jeb.022269

Lentink, D., Dickson, W. B., van Leeuwen, J. L., and Dickinson, M. H. (2009). Leading-edge vortices elevate lift of autorotating plant seeds. Science (New York, N.Y.), 324(5933), 1438–40. doi:10.1126/science.1174196

Lentink, D., Mller, U. K., Stamhuis, E. J., de Kat, R., van Gestel, W., Veldhuis, L. L. M., - van Leeuwen, J. L. (2007). How swifts control their glide performance with morphing wings. Nature, 446(7139), 1082–1085. doi:10.1038/nature05733

Lu, Y., Shen, G. X., and Lai, G. J. (2006). Dual leading-edge vortices on flapping wings. The Journal of Experimental Biology, 209(Pt 24), 5005–5016. doi:10.1242/jeb.02614

Lu, Y., and Shen, G. X. (2008). Three-dimensional flow structures and evolution of the leading-edge vortices on a flapping wing. The Journal of Experimental Biology, 211(Pt 8), 1221–30. doi:10.1242/jeb.010652

Maxworthy, T., (2007). The formation and maintenance of a leading-edge vortex during the forward motion of an animal wing. Journal of Fluid Mechanics, 587, 471–475.

Miau, J. J., Kuo, K. T., Liu, W. H., Hsieh, S. J., and Chou, J. H. (1995). Flow Developments Above 50-Deg Sweep Delta Wings with Different Leading-Edge Profiles. Journal of Aircraft, 32(4), 787–794.

Muijres, F. T., Johansson, L. C., Barfield, R., Wolf, M., Spedding, G. R., and Hedenström, A. (2008). Leading-edge vortex improves lift in slow-flying bats. Science (New York, N.Y.), 319(5867), 1250–3. doi:10.1126/science.1153019

Ol, M. V., and Gharib, M. (2003). Leading-Edge Vortex Structure of Nonslender Delta Wings at Low Reynolds Number. AIAA Journal, 41(1), 16–26. doi:10.2514/2.1930

Ozen, C. a., and Rockwell, D. (2011). Flow structure on a rotating plate. Experiments in Fluids, 52(1), 207–223. doi:10.1007/s00348-011-1215-y

Polhamus, E. C. (1966). A concept of the vortex lift of sharp-edge delta wings based on a leading-edge-suction analogy. National Aeronautics and Space Administration, Nasa Techn. Retrieved from http://hdl.handle.net/2060/19680022518

Shyy, W., Aono, H., Chimakurthi, S. K., Trizila, P., Kang, C.K., Cesnik, C. E. S., and Liu, H. (2010). Recent progress in flapping wing aerodynamics and aeroelasticity. Progress in Aerospace Sciences, 46(7), 284–327. doi:10.1016/j.paerosci.2010.01.001

Srygley, R. B., and Thomas, A. L. R. (2002). Unconventional lift-generating mechanisms in free-flying butterflies. Nature, 420(6916), 660–664. doi:10.1038/nature01223

Taylor, G. S., Schnorbus, T., and Gursul, I. (2003). An Investigation of Vortex Flows Over Low Sweep Delta Wings. 33rd AIAA Fluid Dynamics Conference and Exhibit, 1–13.

Taylor, G., Wang, Z., Vardaki, E., and Gursul, I. (2007). Lift Enhancement over Flexible Nonslender Delta Wings. AIAA Journal, 45(12), 2979?2993. doi:10.2514/1.31308

Usherwood, J. R., and Ellington, C. P. (2002). The aerodynamics of revolving wings I. Model hawkmoth wings. The Journal of Experimental Biology, 205, 1547–1564.

Verhaagen, N. G. (2012). Leading-Edge Radius Effects on Aerodynamic Characteristics of 50-Degree Delta Wings. J. Aircraft, 49(2), 521–531.

Videler, J. J., Stamhuis, E. J., and Povel, G. D. E. (2004). Leading-Edge Vortex Lifts Swifts. Science, New Series, 306(5703), 1960–1962.

Viola, I. M., Bartesaghi, S., Van-Renterghem, T., and Ponzini, R. (2014). Detached Eddy Simulation of a sailing yacht. Ocean Engineering, 90, 93–103. doi:10.1016/j.oceaneng.2014.07.019

Visbal, M. R. (1996). Computed unsteady structure of spiral vortex breakdown on delta wings. AIAA Fluid Dynamics Conference, 2074, 96–2074

Wang, J., Zhao, X., Liu, W., and Tu, J. (2007). Experimental investigation on flow structures over nonslender delta wings at low Reynolds numbers. Journal of Experiments in Fluid Mechanics, 21(2), 1–7.

Wojcik, C. J., and Buchholz, J. H. J. (2014). Parameter Variation and the Leading-Edge Vortex of a Rotating Flat Plate. AIAA Journal, 52(2), 348–357. doi:10.2514/1.J052381

